# Reduced injury risk links sociality to survival in a group-living primate

**DOI:** 10.1101/2022.04.05.487140

**Authors:** Melissa A. Pavez-Fox, Clare M. Kimock, Nahiri Rivera-Barreto, Josue E. Negron-Del Valle, Daniel Phillips, Angelina Ruiz-Lambides, Noah Snyder-Mackler, James P. Higham, Erin R. Siracusa, Lauren J.N. Brent

## Abstract

Affiliative social relationships and high social status predict longer lifespans in many mammal species, including humans. Yet, the mechanisms by which these components of sociality influence survival are still largely unknown. Using 10 years of data and over 1000 recorded injuries from a free-ranging population of rhesus macaques (*Macaca mulatta*), we investigated two possible mechanisms that could underpin the relationship between sociality and survival: sociality (1) reduces injury risk; and/or (2) increases the probability of survival after an injury. We found that sociality can affect an individual’s survival by influencing their risk of injury, but had no effect on the probability of injured individuals dying. Individuals with more affiliative partners experienced fewer injuries compared to less socially integrated. Social status was also associated with lower risk of injury, particularly for older high-ranking individuals. These results represent the first demonstration of a link between social integration and fatal injury risk in a group-living species, and are the first to link social status, injury risk and survival outside of humans. Collectively, our results offer insights into a mechanism that can mediate the well-known benefits of sociality on an individual’s fitness.

---

Uncovering the means by which sociality influences lifespan is of major interest to evolutionary biologists, social scientists and biomedical researchers [1, 2, 3]. Evidence from humans and other animals has provided increasing support for the benefits of affiliative social interactions on survival. The strength of social bonds [4, 5, 6, 7], the number of weak connections [8], the number of associates [9, 10, 11], the number of relatives in a group [12] and the number of indirect connections [11, 13] predict the lifespan of individuals; the general pattern being that those with more social partners are the ones that live longer. Similarly, socioeconomic status in humans and social status in other animals are also robust predictors of mortality risk [14, 15, 12, 5, 16, 17] with lower status individuals suffering a greater risk of death. But precisely how the social environment affects survival is less well understood.

One way for sociality to influence survival is by mitigating the costs of contest competition over resources. Dominance hierarchies, for instance, are believed to have evolved to reduce direct costs associated with competition for resources [18]. Nevertheless, social hierarchies still usually entail disparities in access to resources, with individuals higher in the hierarchy having priority access to food and mates at the expense of their subordinates [19], who may still need to compete for access. Affiliative partners can also help to reduce engagement in agonistic encounters by providing access to resources via cooperation and social tolerance [20]. For example, food sharing, cooperative feeding and co-feeding have been described in several mammals, including some species of bats [21], cetaceans [22, 23, 24], monkeys [25, 26] and apes [20]. Affiliative partners can also help to deter physical aggression from conspecifics by providing agonistic support. For instance, affiliative interactions predict the formation of coalitions in male and female African wild dogs (*Lycaon pictus*)[27], Camargue horses (*Equus caballus*) [28], macaques (*Macaca spp*.)[29, 30] *and chimpanzees (Pan troglodytes*) [31]. Agonistic support has also been widely documented in female-philopatric primate species where related females defend one another [32, 33, 34]. If social status or affiliative relationships reduce the chance of aggressive interactions, these components of sociality may directly enhance survival by allowing individuals to avoid costly outcomes, such as injuries.

In addition to mitigating the immediate costs of aggressive behaviors, sociality may also enhance survival through buffering mechanisms that influence an individual’s health. Differences in access to resources according to social status, for instance, may determine the general body condition and health of individuals. Low social status has been related to higher disease risk [15], higher levels of inflammation [35, 36], reduced healing capacity [37] and overall impaired health in several mammal species, including humans [38, 39, 3]. Affiliative partners, on the other hand, can be valuable resources that can contribute to better health by providing access to food [40, 41] and reducing the burden of infections via hygienic behaviors (*i*.*e*., grooming) [42, 43]. Better health status for high ranking or socially integrated individuals may translate into higher chances of survival in the face of adversity, for example, by improving the chances of healing following an injury.

Yet despite clear hypotheses for the potential mechanisms by which social status and affiliative relationships influence lifespan, there remains a lack of empirical evidence for these mechanisms affecting survival. For example, several studies have shown associations between individual variation in sociality with markers of health and immunity [44, 45, 35, 36], yet the consequences of such differences in the face of naturally occurring challenges to health, and the downstream impact those differences might have on survival are unknown. Similarly, studies supporting a relationship between sociality and lifespan usually do not have the detailed physiological or health data required to test potential mechanisms connecting the two [1]. To fill this gap, we use a long-term data set containing both survival data and detailed information on injuries in a free-living population of rhesus macaques to test whether sociality mitigates the costs of competition (*i*.*e*., injuries) and its consequence on survival.

We explore two injury-related mechanisms that can link sociality with survival. Specifically, we test whether social status and/or affiliative relationships: 1) influence the risk of being injured, and/or 2) alter an individual’s survival trajectory after an injury (Fig. 1). We did so using 10-years of injury data collected *ad-libitum* together with demographic information from male and female rhesus macaques aged 4-29 years living on Cayo Santiago island, Puerto Rico. Rhesus macaques live in multi-male multi-female despotic societies, where access to resources is highly determined by an individual’s position in the dominance hierarchy [46]. Previous studies have shown the benefits of affiliative partners and social status on the survival probability of monkeys in this population [12, 14, 8].Predators are absent from the island, ensuring injuries are mostly the result of physical aggression between conspecifics. Rhesus macaques are seasonal breeders with a mating season that can last from 3 to 6 months. During these periods both affiliative and agonistic interactions are usually heightened [47, 48] and, thus, important trade-offs between health, reproduction and survival may occur [49, 50].

**Figure 1:**
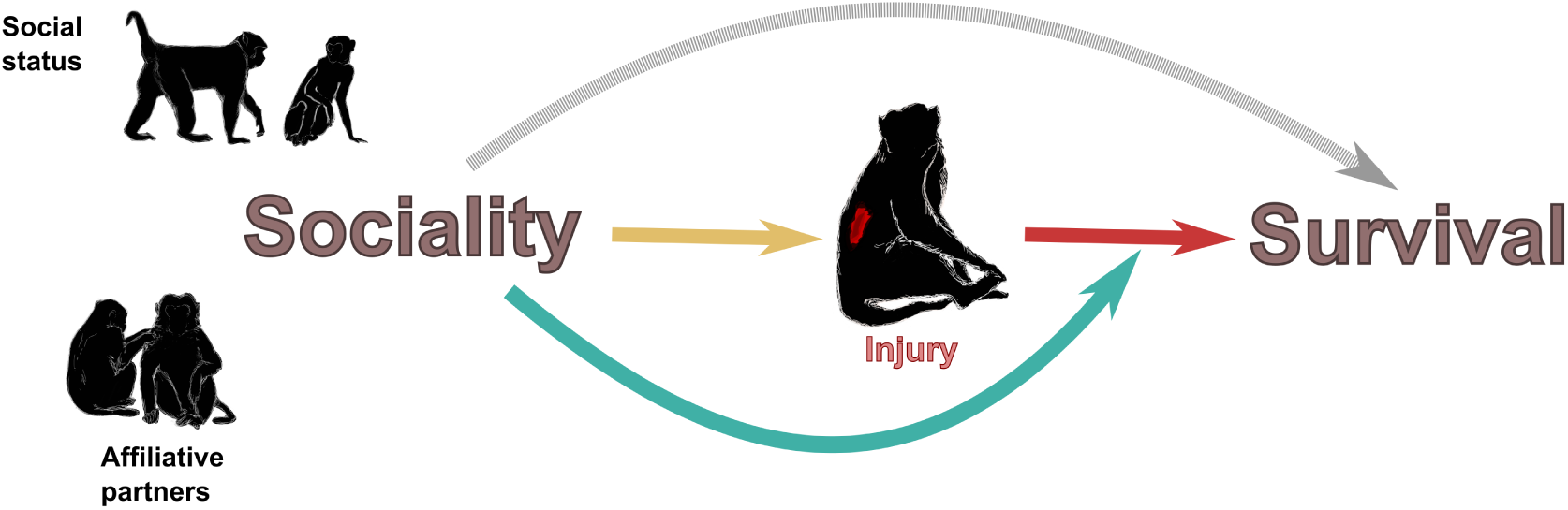
Injury-related mechanisms by which components of sociality (social status, affiliative partners) can influence survival. A direct effect of sociality on survival (gray arrow) has been well established in mammals [4, 5, 7, 10, 11], including studies in the Cayo Santiago population [12, 8]. We explore mechanisms related to injury by which the relationship between sociality and survival might come about. According to the first mechanism, sociality influences the risk of injury (**yellow arrow**) and, therefore, survival (**red arrow**). According to the second mechanism (**green arrow**), sociality affects the survival trajectories of injured individuals.

Because our study hinged on the assumption that being injured was detrimental for survival in this population we first tested whether injuries inflicted by conspecifics increased the probability of death in these animals (Fig 1; red arrow). To test if sociality influences the risk of injury (mechanism 1), we asked whether social status and the number of affiliative partners were associated with an individual’s injury risk (Fig 1; yellow arrow). Given the protective role of high social status and importance of affiliative partners in deterring aggression [20, 32, 18], we predicted that high status individuals and those with more affiliative partners would have a lower risk of injury. To test if sociality can alter the impact of injuries on survival (mechanism 2), we asked if social status and the number of affiliative partners affected the survival trajectories of injured individuals (Fig 1; green arrow). As both social status and social integration can determine differences in health status that may affect healing rates [39, 37, 51], we predicted that high status animals and those with more affiliative partners would have a lower hazard of death from an injury than low status individuals or those with fewer affiliative partners. Our results demonstrate that sociality plays an important role in mediating the risk of injury, offering one of the few clear mechanistic links between sociality and survival in a non-human mammal to date.

## Results

### Effect of injuries on survival

To quantify the extent to which injuries affect an individual’s survival we used time-dependent mixed effects cox models [52, 53]. Animals that were injured were nearly three times more likely to die in the two months following the injury compared to animals that were not injured (Fig. 2A; Hazard (Hz) = 1.06 *±* 0.17 (SEM), z = 6.58, p *<* 0.01, injuries (i) = 1041, deaths (d) = 443, N injured = 571, N uninjured = 1030), independent of their sex or the reproductive season when the injury occurred. Individuals that were severely injured (*e*.*g*. broken bones, exposed organs, multiple wounds or wounds in vital areas, see SI Materials and Methods for details) experienced even a higher hazard of death that was dependent on sex (Hz severity*sexM = 1.46 *±* 0.72, z = 2.02, p = 0.04, i = 398, d = 107, N severely injured = 295). In males, severe injuries were associated with a higher chances of dying compared to non-severe injuries, while in females, severe and non-severe injuries had similar hazards of death (Fig. 2B).

**Figure 2:**
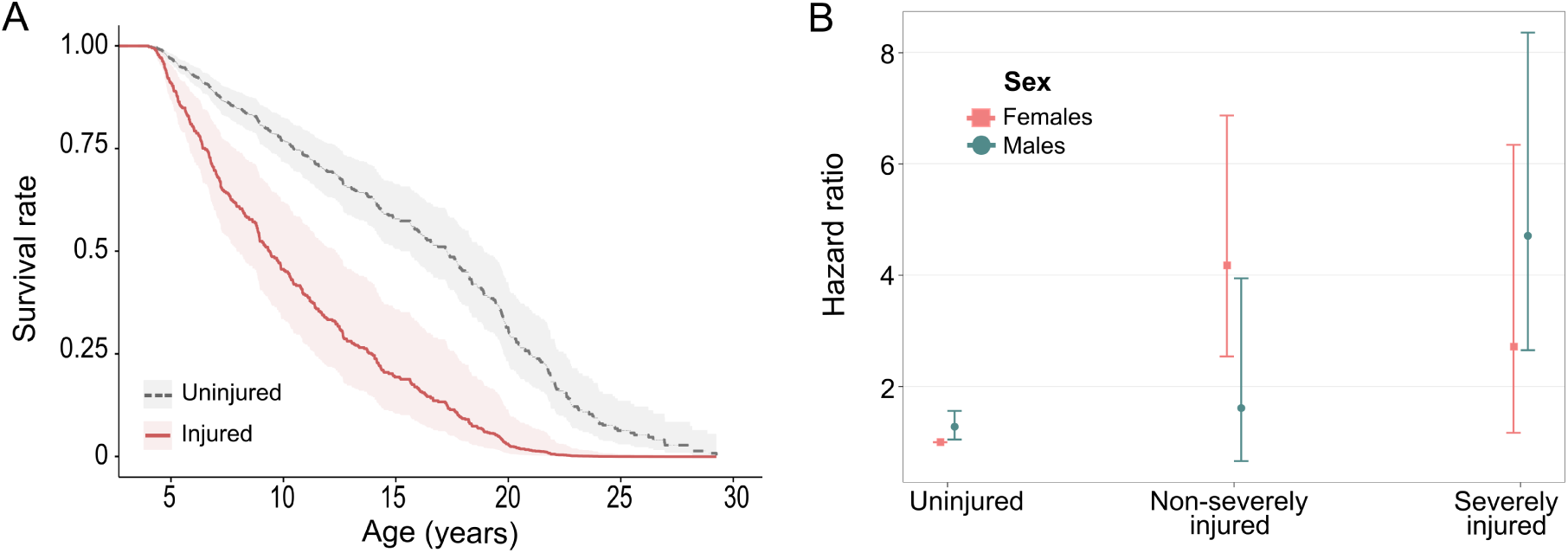
Effect of injuries on survival. **A)** Survival curves adjusted for covariates for injured and uninjured individuals. Injured individuals (red solid line, n = 571, 294 females, 277 males) had near a 3-fold increase in the probability of dying compared to uninjured animals (gray dashed line, *n* = 1030, 557 females, 473 males)(Hz = 1.06*±*0.17, z = 6.58, p *<* 0.01, injuries (i) = 1041, deaths (d) = 443). Curves represent males during the mating season, but those for females were similar. Shaded areas represent standard errors. **B)** Hazard ratios of death for females and males as a function of the severity of injuries. Severe injuries increased the hazard of death relative to non-severe injuries in males (green circles, n uninjured = 473, n non-severely injured = 189, n severely injured = 251), but not in females (Pink squares, *n* uninjured = 557, *n* non-severely injured = 232, n severely injured = 147) (Hz severity*sexM = 1.46 *±* 0.72, z = 2.02, p = 0.04, i = 398, d = 107).

### Mechanism #1: Sociality affects the risk of injury

#### Effect of social status on injury risk

To test if high status animals were less likely to be injured or severely injured than low status ones, we compared their injury risk separately for males and females using logistic models. Given that observations of social interactions were only available for a subset of our subjects, we used proxies of social status previously used in female (matrilineal rank) [14, 12] and male rhesus macaques (group tenure length) [54, 55, 56] to maximize our statistical power. We found that social status in females had a strong effect on the likelihood of being injured, which was dependent on an individual’s age (Odds rankLow*age = 0.3 *±* 0.1, z = 3.02, p *<* 0.01, i = 448, N = 827). Low status females had a higher probability of being injured than high status females, and this probability increased with a female’s age (Fig. 3A). Social status had no relationship with the risk of severe injuries in females (Odds = 0.13 *±* 0.2, z = 0.65, p = 0.5, i = 135, N severely injured = 114). In males, social status also had a strong effect on the probability of being injured which was dependent on age (Odds status*age = 0.1 *±* 0.03, z = 3.28, p *<* 0.01, i = 536, N = 748). In younger males, lower social status was associated with a higher incidence of injuries, while at older ages high status males had higher probability of being injured (Fig. 3B). The same pattern was observed when we focused our analysis on severe injuries (Fig. S2A, Odds status*age = 0.12 *±* 0.04, z = 2.67, p *<* 0.01, i = 245, N severely injured = 168). Consistent with heightened male-male competition over females [48] and with male harassment of females during the reproductive season [57], we also found that injury-risk increased for both males and females during the mating period compared to outside it, independent of their social status (injury: Odds females = 0.85 *±* 0.28, z = 3.02, p *<* 0.01; Odds males = 1.2 *±* 0.26, z = 4.6, p *<* 0.01; severe injury : Odds females = 1.04 *±* 0.26, z = 4, p *<* 0.01; Odds males = 1.38 *±* 0.25, z = 5.4, p *<* 0.01).

**Figure 3:**
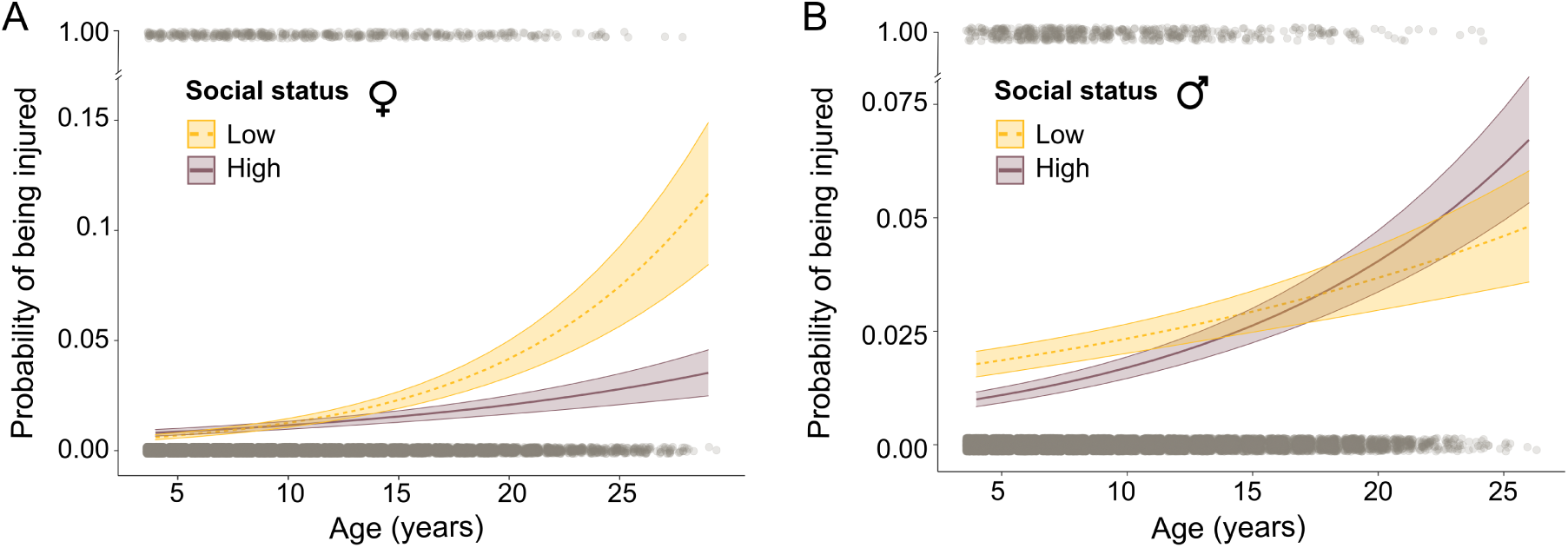
Predicted injury risk in relation to social status. **A)** Injury risk for females as a function of social status and age. Low status females (yellow dashed line, *n* = 420, 237 injuries) had higher chances of being injured than high status females (purple solid line, *n* = 407, 211 injuries), with increasing probabilities for older females (Odds rankLow*age = 0.3 *±* 0.1, z = 3.02, p *<* 0.01). **B)** Injury risk for males as a function of social status and age. For visualization, social status was categorized by selecting the 20th (273 days of tenure) and 80th (2029 days of tenure) percentiles depicting low status (yellow dashed line) and high status (purple solid line), respectively (*n* = 748, 536 injuries). Younger males from low status had higher injury risk than high status young males, yet the opposite occurred at later ages (Odds tenure*age = 0.1 *±* 0.03, z = 3.28, p *<* 0.01). In both plots, shaded areas represent standard errors and gray dots the raw data used in the models (top: injured, bottom: uninjured).

#### Effect of affiliative partners on injury risk

To test whether animals with more affiliative partners were less likely to be injured or severely injured than those with fewer affiliative partners we used logistic models. To support robust statistical analyses, we relied on a proxy (*i*.*e*., number of female relatives in the group) that has been previously shown to influence survival in this population [12]. Female rhesus macaques have a strong bias toward forming partnerships with their maternal kin [58] and this proxy has been positively correlated with network measures of social integration [59]. Males, on the other hand, are the dispersing sex and have few kin in their new groups, and so were excluded from this analysis. We found that the number of close relatives (relatedness coefficient (*r*) = 0.5, *i*.*e*.., mother-daughters and full siblings) in a female’s group had a weak, but not significant, effect on her probability of being injured (Odds = −0.1 *±* 0.05, z = −1.84, p = 0.06, i = 491, N = 851). However, the size of a female’s extended family (r *≥* 0.125, *i*.*e*., spanning three generations) was strongly associated with the likelihood of injury, with females experiencing a 13% reduction in the incidence of injuries for every one standard-deviation increase in their number of female relatives (Fig. 4A; Odds = −0.14 *±* 0.06, z = −2.5, p = 0.01, i = 491, N = 851). The incidence of severe injuries was not affected by the number of close relatives(Odds = −0.06 *±* 0.09, z = −0.6, p = 0.53, i = 147, N severely injured = 123) nor by the size of a female’s extended family (Odds = −0.13 *±* 0.09, z = −1.36, p = 0.18, i = 147, N severely injured = 123).

**Figure 4:**
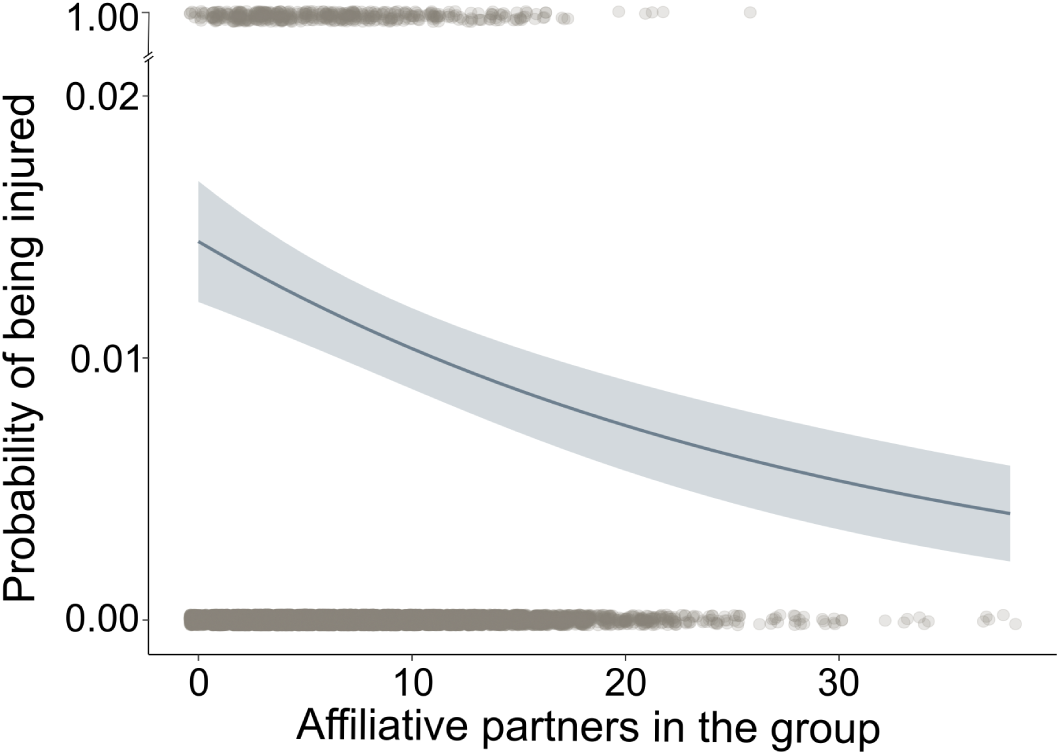
Predicted injury risk as a function of the number affiliative partners. X-axis represents the number of female relatives (extended family, r *≥* 0.125) present in a female’s group (*n* = 851, injuries (i) = 491). Females with more relatives had lower chances of suffering from an injury compared to females with fewer relatives (Odds = −0.14 *±* 0.06, z = −2.5, p = 0.01, i = 491). Shaded areas represent standard errors and gray dots the raw data used in the models (top: injured, bottom: uninjured).

### Mechanism #2: Sociality influences the survival of injured animals

#### Effect of social status on survival of injured animals

To assess whether social status or affiliative relationship buffer the detrimental effect of injuries on survival, we used time-dependent mixed effects cox models. We found no evidence of a buffering effect of social status on the survival of injured females (Hz injured*rankLow = −0.45 *±* 0.46, z = −0.98, p = 0.33, i = 448, d = 103, N = 278) or injured males (Hz injured*tenure = 0.00009 *±* 0.0002, z = 0.47, p = 0.64, i = 536, d = 97, N = 272). Similarly, no buffering effect of social status on survival was observed in severely injured females (Hz injured*rankLow = −0.51 *±* 0.92, z = −0.55, p = 0.58, i = 135, d = 42, N severely injured = 114) or males (Hz injured*status = −0.0001 *±* 0.0002, z = −0.67, p = 0.5, i = 245, d = 57, N severely injured = 168).

#### Effect of affiliative partners on survival of injured animals

We found no evidence for a relationship between survival after an injury and the number of close relatives a female had available at the time (Hz injured*nkin = −0.22 *±* 0.28, z = −0.78, p = 0.43, i = 491, d = 114, N = 294) or current size of her extended family (Hz injured*nkin = 0.03 *±* 0.04, z = 0.59, p = 0.56, i = 491, d = 114, N = 294). Similarly, the number of affiliative partners did not influence the survival of severely injured females (Hz close kin = −0.82 *±* 0.68, z = −1.21, p = 0.23; Hz extended family = 0.008 *±* 0.09, z = −0.1, p = 0.92; i = 147, d = 45, N severely injured = 123).

#### Post hoc mediation analysis

Mediation analyses can be used to test the significance of a mediator in the relationship between an independent and a dependent variable and to measure the effect size of that relationship [60]. Although useful, current mediation analysis approaches are unable to estimate effect sizes for data structured in a logistic manner, such as ours, nor are they able to cope with interaction terms in logistic regressions [61]. Given these limitations, we could not use mediation analysis to evaluate if the effect of social status on survival was mediated by injury risk because these results relied on an interaction with individual age (Fig. 3), nor could we use it to estimate the effect size of any of our results. We did, however, use mediation analysis to assess if injury risk significantly mediated some of the effect of affiliative partners on survival. Our mediation analysis confirmed a direct effect of affiliative partners on survival by showing that the size of a female’s extended family significantly reduced her hazard of death (direct effect = −0.065 *±* 0.02, z = −3.19, p *<* 0.01). It also confirmed that this relationship was significantly mediated by the risk of being injured (indirect effect z-score = −2.31, p *<* 0. 05).

## Discussion

Taken together, our results suggest that different components of the social environment can modulate the risk of suffering an injury and, therefore, the hazard of death. We found that high social status was associated with a lower injury risk for specific periods of males’ and females’ lives, and that a female’s number of affiliative partners may help to prevent injuries. In contrast to previous research showing that individuals with higher social status had faster healing rates [37], we found that none of the measures of sociality analyzed affected the survival trajectories of injured animals.

To our knowledge, this is the first field study to quantify the consequences of injuries on the probability of death in a nonhuman primate. Other studies in wild populations of baboons (*Papio sp*.) and Afro-eurasian monkeys have established the social and demographic predictors of injury risk [62, 37, 63, 64, 65, 66], yet its consequences on survival have yet to be shown. We found sex-differences on the influence of severe injuries in survival that can reflect trade-offs between the energy allocated for reproduction versus immunity [4, 37].For instance, during the reproductive season the probability of being severely injured was substantially higher for both sexes. During this period, males may be particularly immunocompromised given the high amount of energy and resources required to sustain the effort associated with mating [49, 67], which can impair injury recovery. On the other hand, females usually have higher demand on their immune systems during lactation [4, 68], *i*.*e*., outside the reproductive season. Therefore, females may cope better than males with severe injuries during the reproductive season at the expense of being more susceptible to the consequences of injury outside this period.

We found support for one of the hypothesized mechanisms linking sociality to survival, whereby sociality reduces an individual’s risk of injury. High social status animals were injured less than those of lower status during specific periods of their life, and females with more affiliative partners (i.e., kin) were less likely to be injured than less integrated females and, thus, experienced lower hazard of death. Our results linking social status to reduced risk of injury are consistent with the skewed access to resources in systems with clear linear dominance hierarchies [19]. High status individuals may not need to engage in costly aggression for food or mates, in contrast to low status animals who must gain access through contests. Although we could not test for a mediation effect of injury risk on the relationship between social status and survival, our results suggest that low status individuals experience greater hazard of death as a result of enhanced risk of injury. Our finding that social status did not influence the risk of injury in young females may be because at younger ages females’ relative positions in the dominance hierarchy have yet to be fully established [69]. Further, we showed that older high status males were more likely to be injured than older low status males. This finding may reflect heightened aggressive challenges from lower status animals to those higher in the hierarchy as a consequence of a decline in the body condition with age [70] and, thus, the capacity of older high status males to maintain their dominance.

Previous studies in matrilineally-structured primate species, in which most of the affiliative relationships are with female relatives, have shown that females commonly engage in agonistic encounters to support and protect their kin [32, 33, 34], even when confronting higher status individuals [71]. In line with these studies, our results suggest that having more relatives available may provide a numerical advantage to deter physical aggression. Other mechanisms, such as social tolerance when accessing resources [20] could also explain fewer injuries in the presence of more affiliative partners. Interestingly, only the size of a female’s extended family, not her number of close relatives, had a significant relationship with risk of injury. This suggests that the number of close relatives in a group (range in our study: 0-5) may not be enough to provide robust agonistic support or access to resources, compared to the size of a female’s extended family (range in our study: 0 - 38).

We found no support for the second hypothesized mechanism that we explored to link sociality to survival; none of the measures of sociality analyzed influenced an individual’s survival trajectory following injury. Despite a vast body of literature supporting differences in health and immunity between individuals of different social status [44, 35, 39], we found no evidence for an effect of social status on the survival trajectories of injured animals. These findings contrast with a previous study on wild baboons where high status males had faster healing rates than lower status males [37]. Although we did not quantify differences in healing times, our results suggest that the probability of recovering from an injury was not influenced by an animal’s position in the dominance hierarchy. These differences might be explained in part by differences in features of the two study systems. Animals on Cayo Santiago are provisioned with food on a daily basis and access to the nutrients needed to support immune function might not be as skewed as they are in the wild [45].Notwithstanding, in both systems high social status has been associated with elevated levels of glucocorticoids and androgens [72, 73, 50], well known immune-suppressors, which suggest that in the Cayo Santiago population, unlike the baboons, the benefits of being of high status may not outweigh the costs in terms of injury recovery.

We also found, contrary to our predictions, that the benefits associated with affiliative partners, such as feeding tolerance [74, 75] and social hygienic behaviours [42, 43], seem not to have helped females to cope with the detrimental effect of injuries on their survival. It is possible that social hygienic behavior, such as removal of ectoparasites by grooming, have long-term health benefits but do nothing to enhance the short-term immune response required to heal damaged tissue [76].Additionally, grooming wounded areas may, in fact, be detrimental for the healing process as it could lead to the removal of protective scabs [43]. This could be one reason why females with more affiliative partners, who are presumed to receive more grooming and to have more access to food via social tolerance, did not have improved survival trajectories after an injury. Previous research on this population has shown that the number of close relatives and the size of a female’s extended family are associated with increased survival probability [12, 8]. The results of the current study suggest this relationship does not come about because of the reduced risk of death from injury. Further research is needed to elucidate to what extent other mechanisms involving health differences (e.g., disease susceptibility) play a role in the benefits of social partners in the survival of females in this population. Additionally, direct behavioral observations in a large sample of individuals with paired injury data will be required to explore refined ego-networks characteristics and to expand these results to affiliative relationships of males and unrelated females.

In sum, our study provides evidence for a mechanism linking sociality to lifespan. Growing literature has supported a strong relationship between the social environment and survival in many mammal species [3], but the ultimate function of some components of sociality, such as social relationships, remain unclear [77]. Although sociality has been demonstrated to enhance health and immunity [44, 35, 45], here we showed that these benefits did not translate to an improved ability to cope with the risk of death from injuries. Instead, we found that sociality plays an important role in preventing individuals from suffering injuries that would likely lead to death. Given how rare injuries are in this population, we do not expect that this is the only mechanism linking sociality to survival. Other mechanisms may include sociality-mediated differences in components of health related to disease susceptibility. In wild animal populations, social partners may also help with predator detection [78], predator mobbing [79], finding food sources [80], thermoregulation [81], among other possibilities. Nevertheless, here we provide rare empirical evidence for an ultimate function of social relationships, showing one mechanism by which high status and socially integrated individuals live longer. Demonstrating the relative importance of different mechanisms linking sociality and survival will be challenging but a crucial goal of future research. Our study provides insight into the essential role that long-term datasets that combine both demographic and health data will play in meeting this challenge.

## Materials and Methods

### Subjects

We studied a population of free-ranging rhesus macaques on the island of Cayo Santiago in Puerto Rico. The island is home to a population of *∼* 1800 individuals living in 6-10 mixed sex naturally formed social groups. The field station is managed by the Caribbean Primate Research Center (CPRC), who monitor the population daily, and maintain the long-term (*>*75 years) demographic database including data on births, deaths, social group membership for all animals and a genetic parentage database for animals born after 1992 [82]. Animals have *ad-libitum* access to food and water, the island is predator-free and there is no regular medical intervention for sick or wounded individuals. We focused on all subadult and adult females and males between 4 and 29 years of age that were alive between the years 2010 and 2020, a period for which records on injuries exist (see below for details on how injury data was collected). In this study we included data on 571 injured individuals (294 females, 277 males) and 1030 uninjured individuals (557 females, 473 males). From these animals, 342 (85 injured, 258 uninjured) were removed from the population by the CPRC for population control purposes [83]. For all individuals, birth dates were known within a few days. Removal dates were known for all removed individuals. Dispersal from the island almost never occurs, therefore death dates were also known within a precision of a few days.

### Observation of injuries

From 2010 to 2020 CPRC staff collected *ad-libitum* observations on the incidence and recovery of injuries, during the daily monitoring of social groups for demographic purposes. Monkeys were individually recognized based on their identity tattoos located on their chest and leg. Whenever a staff member noticed a wounded animal or an animal displaying signs of injury (*e*.*g*. bleeding, limping), they recorded the animal ID, type of injury and additional details on the general state of the animal (*e*.*g*. by evidence of weight loss or poor physical condition). If there was a visible wound, observers additionally recorded the area of the body affected, if it was a recent or old wound based on the presence of scars, and whenever possible, an estimate of its size. Observers updated the records every time they encountered the injured animal during their daily census routine with an average update time for an injured individual across the 10 years of 42.17 days. In total, 1137 injury events were observed with an average of 107.6 ± 63.5 per year. Here, we included all the records of injuries that were considered non-ambiguous (*i*.*e*., those with visible damage to the skin) including bites, scratches, cuts and abrasions along with other clearly observable injuries such as fractures and exposed organs. Our final sample consisted of 1041 injuries collected from September 2010 to April 2020. We classified these injuries based on their degree of severity, where severe injuries were those involving broken bones, exposed organs, multiple wounds and any wound in vital areas, including head, neck, abdomen or genitalia (*n* = 398). All other injuries were considered non-severe (*n* = 643).

### Measures of sociality

We used proxies of social status (dominance rank) in our analyses. Observations of agonistic interactions between pairs of animals-from which dominance rank is often computed-were only available for a subset of subjects (194 unique individuals injured in 292 injury events, 485 uninjured individuals). To maximize statistical power, we decided to use the complete dataset and to use known proxies of social status instead; group tenure in males [54, 55, 56] and matrilineal rank in females [14, 12]. Male rhesus macaques reach dominance through queuing [84]; those that have been in a group for longer are usually high-ranking [54]. We determined tenure length using information on monthly social group membership. Group tenure length was computed as the time (in days) a male has been observed in his current group at the date of interest (current date minus date of dispersal). If a male had not yet dispersed and remained in his natal group, we computed group tenure since their birth date. If a male died or was removed from the population before the end of the period of interest, we computed group tenure up to that point. We established tenure length for all the males in our dataset (*n* = 750, *n* injuries = 550). However, 67 of those males had periods where they were observed living outside a social group (*i*.*e*., they were “extra-group”). These specific periods when group tenure could not be computed were dealt differently depending on the analysis in question and we discuss this on a case-by-case basis below.

Female rhesus macaques are philopatric and form maternally inherited stable linear dominance hierarchies whereby daughters occupy a rank just below their mothers [85]. Members of a same matriline tend to be adjacent to one another in the hierarchy, thus the rank of an entire matriline can be used as a proxy for individual rank in social groups containing more than one matriline [14]. We determined matrilineal rank using known social status based on pairwise agonistic interactions from females in our dataset. We identified only one matriline per group as ‘high-ranking’ - the one containing the alpha female - while all the others in the group were classed as ‘low-ranking’. Females in groups with a single matriline were disregarded as rank is a relative measure and females from groups with a single matriline are all of the same rank. We established matrilineal rank for 827 females (407 high ranking, 420 low ranking, *n* injuries = 448).

To confirm that group tenure and matrilineal rank were appropriate proxies for social status we looked at the correlation between dominance rank computed from animals with known social status based on agonistic interactions and our proxies. The correlation between group tenure and dominance rank-measured as the percentage of same-sex animals outranked in the group [86] - was moderate and significant (Fig. S1A; *Pearson’s r* = 0.62, p *<* 0.01). Matrilineal rank and categorical dominance rank were strongly correlated (high-ranking: *≥* 80% outranked, low ranking: ≤ 79% outranked [87]) based on Cramer’s V coefficient (Fig. S1B; *Cramer’s V* = 0.39, chi-square = 159.42, *p <* 0.01), which measures the association between two categorical variables [88].

As above, we only had data on affiliative interactions for a subset of our subjects. Therefore, to maximize our sample size we followed a previous study [12] and used the number of female relatives (4 years and older) that were present in a female’s social group as a proxy for social capital. Female rhesus macaques preferentially interact with their female kin compared to non-kin individuals [58], thus those with greater number of relatives are expected to have more opportunities for social support. We limited this approach to females as males, being the dispersing sex, often have very few close kin in their new groups, and might not be able to recognise unfamiliar kin [89]. Using the Cayo genetic pedigree database we computed the number of close kin (r = 0.5) and extended family (r *≥* 0.125) for all injured and uninjured females in our dataset (*n* = 851, *n* injuries = 491). We decided to test these two levels of relatedness as the first represents the strongest kin-bias (*i*.*e*., mother-daughter or full sisters) and the second the lowest threshold for kin bias in affiliative interactions for rhesus macaques [90].

### Statistical approach

For all of the statistical analyses we defined a two-month time window (hereafter, bimonthly interval) as the period from which the injury status could transition from injured to not injured based on the average update time for an injured animal (*i*.*e*., average time between two consecutive records) and the computed average healing time. Thus, all variables were evaluated on a bimonthly basis (*i*.*e*., each row in the dataset represents a two-month interval). For each of the questions we ran two models, one that included injury status based on all injuries (model 1) and other that included injury status for severe injuries only (model 2).

### Effect of injuries on survival

To establish the effect of injuries on survival we used time-dependent Cox proportional hazard (PH) models [52]. For the analyses we used the whole dataset (*n* = 1061), including injured and uninjured animals from both sexes. Animals that were removed from the population or that were still alive at the end of the study period were censored. The predictor of interest was the injury status (*i*.*e*., all injuries or severe injuries) along with other relevant variables that may influence survival probability, such as reproductive season (*i*.*e*., mating vs no-mating) and sex. Age was accounted for implicitly in the models. Additionally, we included random effects for the specific bimonthly interval within the study period to control for potential mortality sources at the population level and individual identity to account for repeated measures. To determine the bimonthly interval we divided the whole study period (10 years) in intervals of two months-ranging from 1 to 58 - where 1 represents the first two months since September 2010. We tested for interaction effects among our predictors and only retained them if statistically significant to avoid issues of overfitting.

### Mechanism #1: Sociality affects the risk of injury

To assess the effect of social status and the number of affiliative partners on the risk of injuries, we used generalized linear mixed models with binomial distribution (logit models). In all the models we asked whether our measures of sociality influenced the probability of being injured in a given bimonthly interval. To test if high status animals were less likely to be injured compared to low status ones, we ran the analyses separately for each sex (*n* females = 827, *n* males = 750). For males, social status was estimated from group tenure computed up to the end of each bimonthly interval. Bimonthly intervals where males were extra-group and so group tenure could not be computed, were excluded. For females, we used matrilineal rank, which remains constant across the lifespan and, thus, remained the same in every interval. To test if animals with more affiliative partners were less likely to be injured compared to animals with social partners we used only females (*n* = 851), fitting separate models for the two thresholds of relatedness (close kin and extended family). The number of relatives present in a group was computed for each bimonthly interval. We modelled injury status as a function of social status or number of affiliative partners, while controlling for age and reproductive season. As group tenure and age could be correlated, we checked for collinearity between these predictors using the variance inflation factor (vif), but no correlation was found (vif = 1.01). Random effects were included for individual ID - to account for repeated measures - and for the specific bimonthly interval within the study period. We z-scored continuous variables to help convergence and tested interaction terms among all our predictors, which were retained if significant.

### Mechanism #2: Sociality influences the survival of injured animals

To examine the effect of sociality (social status and number of affiliative partners on the survival of injured animals we used time-dependent cox ph models. As before, we tested for an effect of social status on survival in separate models for males and females and examined only females to test the effect of affiliative partners on survival post-injury. In all the models the predictor of interest was specified by an interaction term between injury status and the sociality measure. Variables were evaluated on a bimonthly basis with a time-dependent covariate for reproductive season. Random effects were included for individual ID and bimonthly interval. We additionally included a time-dependent fixed effect for group size to control for its potential effect on the number of kin available and on survival [2]. As some bimonthly intervals had missing information for group tenure, we ran two models for males; a complete case analysis and a model using mean-matching multiple imputation with 20 iterations to fill the missing data [91, 92], yet the estimates were identical between both procedures. Given that the main predictor was an interaction term, we did not attempt to fit other interactions.

### Post hoc mediation analysis

To further confirm our findings that sociality significantly influences survival by reducing risk of injury we ran a mediation analysis. Given limitations to use mediation analyses with different type of models (logistic and cox), we translate our cox model to predict survival into a logistic regression, where the outcome represents if the animal was still alive (0) or death (1) as a function of injury status on each bimonthly interval. Unlike cox models, logistic regressions can not handle individuals for which the outcome is unknown (*i*.*e*., censored), therefore for those individuals the last bimonthly interval in the study was not considered. Different methods for testing mediation using logistic models have been proposed. However, to date there are still no robust methods to quantify the effect size or to consider interaction terms [61]. Given this limitation, we were only able to test the significance of the mediation effect of injury risk on the relationship between the number of affiliative partners (r *≥* 0.125) and survival. We ran first a model where the number of affiliative partners and covariates predicts the injury risk (injuries *∼* sociality + covariates). From this model, we extracted the estimate and standard error for affiliative partners. Then, we ran a second model where both sociality and injury risk predict survival (survival *∼* sociality + injuries + covariates), and extract the estimate and standard error for injury risk. Finally, we computed the standardized element (z-score) following Iacobucci [93]. We determined significance by contrasting the z-score against a standard normal distribution, thus an absolute value greater than 1.96 represents a statistically significant mediation effect.

### Ethics

This research complied with protocols approved by the Institutional Animal Care and Use Committee (IACUC) of the University of Puerto Rico (protocol no. A6850108) and by the University of Exeter School of Psychology’s Ethics Committee. The Caribbean Primate Research Center (CPRC) Animal Care and Use Program is evaluated and approved by the IACUC.

## Acknowledgments

We thank the CPRC for the permission to undertake research on Cayo Santiago, along with Edgar Davila, Julio Resto, Bianca Giura and Giselle Caraballo, who assisted in the collection of injury data. Additionally, we thank members of the Brent lab and CRAB for their helpful suggestions, especially Sam Ellis, Delphine De Moor, Michael Weiss, Jordan Hart and Andre Pereira. This work was supported by ANID-Chilean scholarship [number 72190290], the National Institutes of Health [grant R01AG060931 to N.SM., L.J.N.B. and J.P.H., R00AG051764to N.S-M], a European Research Council Consolidator Grant to L.J.N.B [Friend Origins - 864461], a MacCracken Fellowship to C.M.K and a National Science Foundation Doctoral Dissertation Research Improvement Grant to C.M.K. [1919784]. The CPRC is supported by the National Institutes of Health. An Animal and Biological Material Resource Center Grant [P40OD012217] was awarded to the UPR from the Office of Research Infrastructure Programs (ORIP), and a Research Facilities Construction Grant [C06OD026690] was awarded for the renovation of CPRC facilities after Hurricane Maria.

## Notes

### Competing Interest Statement

The authors have declared no competing interest.

### Summary of Updates

Group size results excluded

